# Coordinated network of T cells and antigen presenting cells regulate tolerance to food

**DOI:** 10.1101/2024.07.11.603064

**Authors:** Anna Rudnitsky, Hanna Oh, Joanathan Talmor, Ranit Kedmi

## Abstract

To efficiently absorb nutrients and facilitate microbial commensalism, the host establishes tolerogenic immune programs against dietary and commensal antigens, promoted by peripheral regulatory T cells (pTregs)^1,2^. Previous research into which antigen-presenting cells (APCs) initiate dietary pTreg responses focused on type 1 DCs (cDC1)^3^. However, we now report that food-specific pTreg cells are exclusively induced by the recently identified RORγt+ APCs^4–8^, and not by cDC1. Instead, pTregs interact with cDC1 to regulate the response of food-specific CD8αβ T cells that accumulate in the lamina propria (LP) and epithelial layer of the small intestine (SI) and express memory markers. Upon infection with pathogens that mimic dietary antigens, food-specific CD8αβ cells activate an effector program to potentially guard against ‘Trojan horse’ attacks. Uniquely, after the infection resolves, these cells do not respond to their corresponding dietary antigens, allowing for safe food consumption. Based on our findings, we propose that in response to dietary antigens, dedicated antigen-presenting cells direct a unique CD8αβ response that is coupled to the pTreg program to facilitate protective acute effector responses within the overall strategy of tolerance.

## Main

Oral tolerance is a regulatory immune response to dietary antigens that depends on peripheral regulatory T (Treg) cells ^9,10^ and suppresses inflammation to those same antigens if encountered in an inflammatory setting, thus being essential for survival. Migratory cDC1 are thought to drive the food-specific pTreg response^11,12^, since they present dietary peptides to T cells^13^, foster Treg differentiation in vitro^11^, and are located proximal to primed T cells in the gut-draining mesenteric lymph nodes (MLNs)^14^. Paradoxically, genetic depletion of cDC1s does not result in loss of pTreg or oral tolerance loss ^3,15^. Recently, we found that RORγt+ APCs^4^, and not cDCs, are the exclusive APCs that drive pTreg generation against gut microbial antigens ^4–6^, which prompted us to revisit the cellular requirements for dietary immune responses.

Here we report that food-specific pTreg cells are exclusively induced by RORγt+ APCs which, similar to the response to Hh, utilize integrins αvβ8 to potentially activate TGFβ ^4,5^.While cDC1s are not essential for pTreg induction, they initiate a unique subset of resident and memory-like CD8αβ T cells to food antigens. This food-specific CD8αβ response is regulated by pTregs, which under homeostasis restrict the CD8αβ T cell expansion by attenuation of the co-stimulatory signals on cDC1 cells. Food-specific CD8αβ T cells can acquire an effector program to provide protection against pathogens expressing antigens shared with food. Yet, once the pathogen is cleared, unlike conventional tissue resident memory cells (Trm), these CD8αβ cells regain a tolerance phenotype toward the cognate food antigens. The findings of a hierarchical interaction between RORγt+ APCs, pTregs and cDC1s for maintaining oral tolerance provide new light on a cellular network that prevents inflammatory responses to food antigens while keeping the host poised against microbial infections.

### RORγt+ APCs induce dietary pTreg response

To study the potential role of RORγt+ APCs in directing the differentiation of food-specific pTreg cells we first monitored dietary-specific CD4+ T cells differentiation in mice deficient for antigen presentation by RORγt-lineage cells, including both ILC3 and Janus cells or Thetis cells (*Rorc-Cre;I-Ab f/f*, or MHCII-OFF^ΔRORγt^ mice) ^4–6,8^. We transferred naive Ovalbumin (OVA)-specific CD4+ T cells (OT-II cells isolated from OT-II T cell receptor transgenic mice) into MHCII-OFF^ΔRORγt^ mice orally administered OVA as a food antigen source (Fig. 1a). Differentiation of transferred OT-II cells into pTregs was completely abrogated in MHCII-OFF^ΔRORγt^, but not in control mice (Fig. 1b), indicating that antigen presentation by RORγt+ APCs is required for the induction of food-specific pTreg.

**Figure 1:**
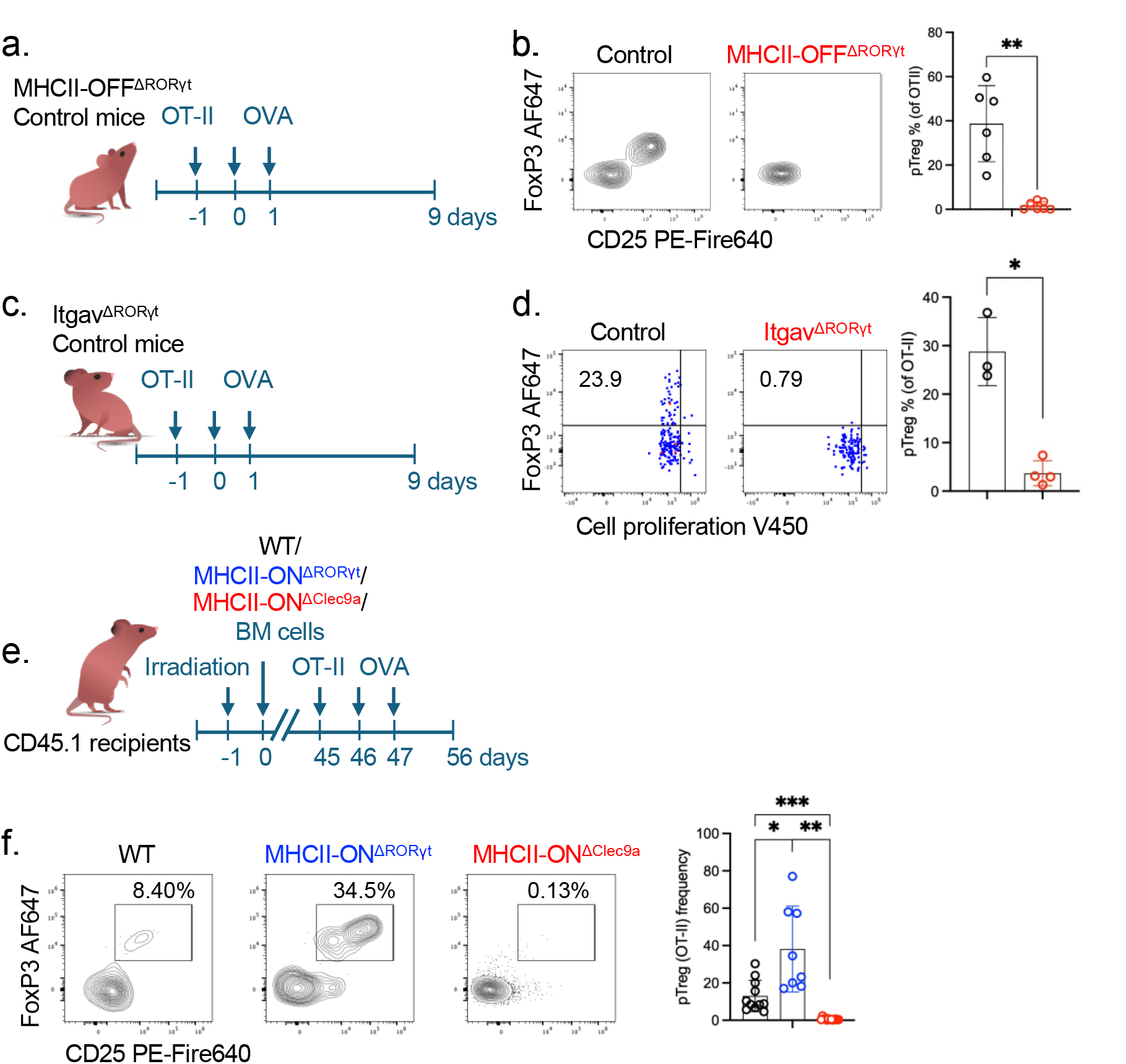
RORγt APCs, but not cDC1, induce a pTreg program in response to dietary antigens. (a) Experimental design of testing food-specific pTreg induction in MHCII-OFF^ΔRORγt^ mice. (b) Representative flow cytometry (left) and aggregate data (right) of OT-II T cell differentiation in MLN of MHCII-OFF^ΔRORγt^ (n = 7) and littermate control (MHCII-/lsl)(n=6) that were orally administrated with OVA, 10 days after transfer of naive TCR transgenic T cells. Data summarize two independent experiments. (c) Schematic of testing food-specific pTreg induction in Itgav^ΔRORγt^ mice. (d) Proliferation and differentiation of OVA-specific pTreg cells in the MLN of Itgav^ΔRORγt^ (n = 3) and littermate control mice (n = 4). V450-labelled OT-II T cells were analyzed at 3 days after adoptive transfer into mice orally administrated with OVA. Representative flow cytometry profiles (left) and aggregate data (right). (e) Experimental design of studying food-specific pTreg induction in BM chimeric mice, reconstitute with alternative BM cells allowing exclusive antigen presentation by APC subsets, as indicated. (f) Representative flow cytometry profiles (left) and aggregate data (right) of OT-II T cell differentiation in MLN of bone marrow chimeric mice reconstituted with donor bone marrow cells as indicated, that were orally administrated with OVA, 10 days after transfer of naive TCR transgenic T cells. wild-type (WT) (n = 11), MHCII-ON^ΔRORγt^ (n = 8) and MHCII-ON^ΔClec9a^ (n = 13). Data summarize three independent experiments. Error bars: means ± s.e.m. Each symbol represents an individual mouse. Statistics were calculated by two-tailed unpaired t-test; *P < 0.05; **P < 0.01; ***P < 0.001.

RORγt+ APCs not only instruct generation of microbe-specific T cells through antigen presentation, but also require expression of integrins αvβ8, which activates TGFβ that contributes to pTreg differentiation^4–6^. Transferred OT-II cells labelled with violet cell proliferation dye (VPD450) exhibited proliferation by day 3 in small intestine-draining mesenteric lymph nodes (MLN) of OVA-fed control mice, with upregulation of FOXP3. By contrast, mice genetically deficient for integrins αv in RORγt+ APCs *Rorc-Cre*;*Itgav f/f* or Itgav ^ΔRORγt^) failed to induce OVA-specific pTreg cell differentiation, pointing to a shared molecular mechanisms for generation of microbial and food pTreg responses (Fig. 1c-d).

As antigen presentation and integrins αvβ8 are required by RORγt+ APCs for the induction of pTreg responses to gut commensals^4,5^, it is expected that MHCII-OFF^ΔRORγt^ and Itgav ^ΔRORγt^ mice may suffer from gut dysbiosis that could indirectly impact the generation of food-specific pTreg responses. We tested reciprocally whether antigen presentation limited only to RORγt-lineage cells is sufficient to enable food-specific pTreg cell differentiation. We reconstituted irradiated CD45.1 mice with bone marrow from mice expressing MHCII only in RORγt+ APCs, and not in other cells (*Rorc-Cre;I-AB* −*/lsl* or MHCII-ON ^ΔRORγt^), or from WT control mice (Fig. 1e). Consistent with our hypothesis, these analyses revealed that antigen presentation by RORγt+ cells alone was sufficient to induce the pTreg response to food (Fig. 1f).

Since the DC subsets were suggested to have redundant functions, we next tested whether cDC1s are additionally capable of promoting pTreg responses alongside RORγt+ APCs. We reconstituted irradiated CD45.1 congenic mice with bone marrow from mice expressing MHCII only by cDC1 and cDC2 but not by RORγt+ APCs (*Clec9a-cre;I-AB* −*/lsl* or MHCII-ON^ΔClec9a^ mice). Remarkably, chimeric mice reconstituted with MHCII-ON^Δ Clec9a^ failed to differentiate transferred OT-II cells into food-specific pTregs (Fig. 1e-f, Extended Data Fig. 1a-b). In fact, in these chimeric mice, OT-II cells acquired a Th1 program (Extended Data Fig. 1c). Notably, chimeric mice reconstituted with a mix of MHCII-ON^ΔRORγt^ and MHCII-ON^ΔClec9a^ donor cells displayed a pTreg response and not a Th1 program (Extended Data Fig. 1c). Collectively, these results suggest that cDCs are unable to induce pTreg responses, and that RORγt+ APCs have a unique role in instructing pTreg generation. Additionally, it implies that, in the absence of pTreg cells, Clec9a-lineage cells, potentially cDC1, promote a Th1 program in response to dietary antigens.

### Food-specific CD8αβ responses in intestinal barrier tissue

Since our data propose a “division of labor” model, we next turned to study the role of the cDC1 subset in generating immune responses to food antigens. While cDC1 have been shown to present food peptides and interact with food-specific CD4 T cells, our data suggest that they do not initiate the pTreg program, which raises questions as to the role of cDC1 in generating immune responses to food antigens. cDC1 are mostly known for activating the Th1 program^16,17^ and CD8 T cells through cross-presentation^18^. The role of CD8 T cells in dietary immune responses is controversial^19^ and was largly neglected following studies that found these cells to be unnecessary for inducing oral tolerance^20–22^. Since we could not identify a Th1 response to dietary antigens at homeostasis, we decided to test whether food induces a diet-specific CD8αβ response. Three days following oral exposure to OVA, transferred OVA-specific CD8+ T cells (OT-I cells isolated from OT-I TCR transgenic mice) were proliferating in the MLN (Fig. 2a,b). We could detect them later in the small intestine lamina propria (SILP) and in the intestinal epithelial layer (Fig. 2c-d, Extended Data Fig. 2a,b). OVA-specific CD8αβ T cells persisted in the epithelial layer even in the absence of antigen, and were detectable 60 days following initial administation of OVA (Extended Data Fig. 2d,e). In addition to transferred cells, we were also able to detect endogenous OVA-specific CD8αβ cells in the SILP and in the epithelial layer (Fig. 2e-g). Consistent with their maintenance in the tissue, these cells exhibited a Trm-like phenotype, as they were found to express CD69 and CD103 residency markers, as well as the anergic markers CD73 and FR4, and were negative for the effector marker KLRG1 (Fig. 2h and Extended Data Fig. 2c). This shows that short-term exposure to new food antigens can induce an antigen-specific population of long-lasting intestinal non-effector CD8αβ ‘Trm-like’ cells.

**Figure 2:**
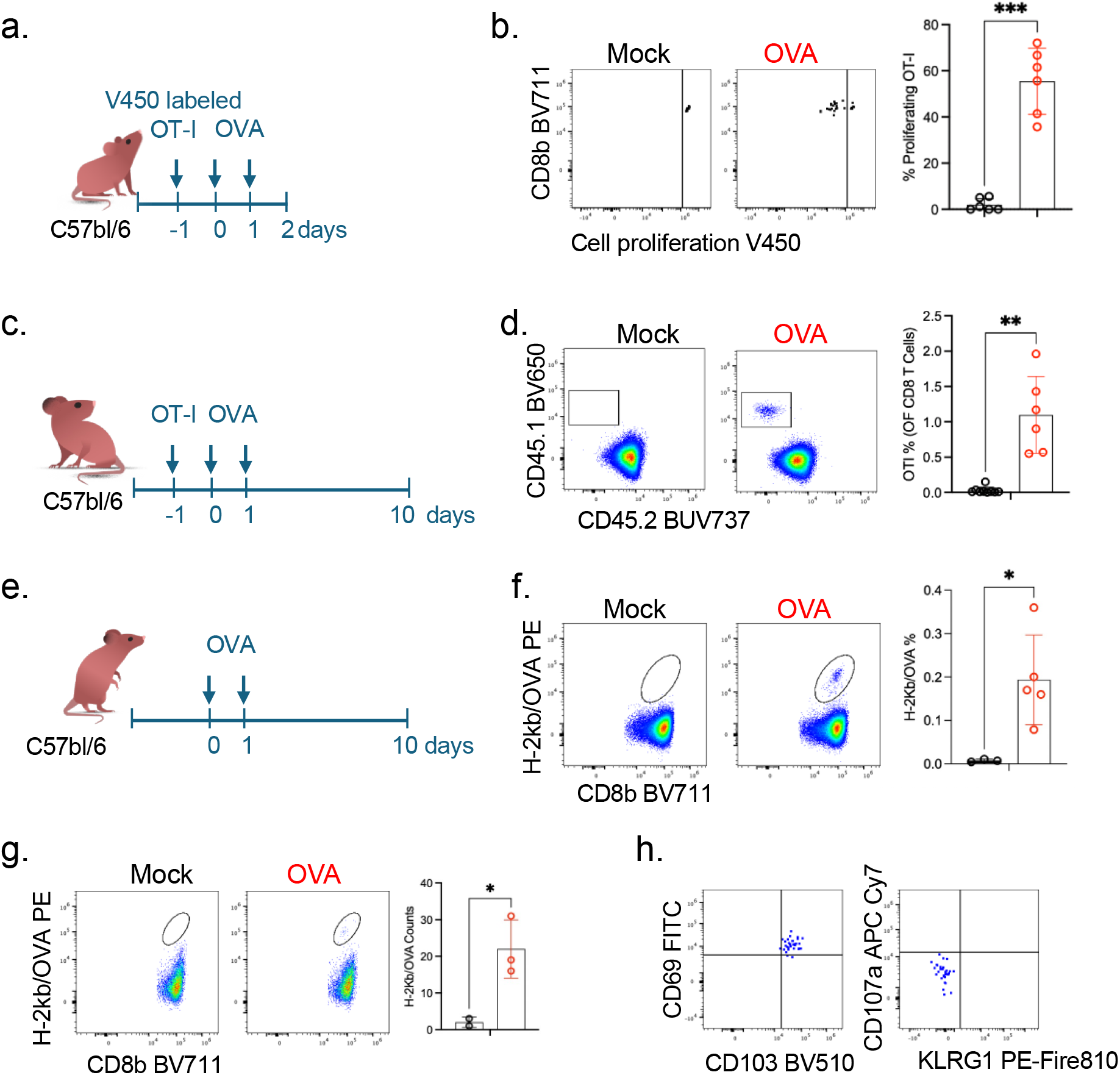
Food-specific ‘Trm-like’ CD8αβ response in the MLN and LP. (a,c,e) Experimental design of testing food-specific CD8 induction in the MLN SILP as indicated. (b,d) Representative flow cytometry profiles (left) and aggregate data (right) of TCRtg OVA-specific CD8 T cell (OT-I) expansion in MLN (b) and SILP (d) in mice orally administrated with OVA (n=6) or PBS (n=6)10 days following OT-I transfer. (f,g) Representative flow cytometry profiles of endogenous OVA-specific CD8 cells in the SILP (f) and in the SI epithelial layer (g) 10 days after mice were orally introduced with OVA (n=5) or PBS (n=3). (h) Representative flow cytometry profiles of CD69, CD103, KLRG1 and CD107a in endogenous OVA-specific CD8 cells in the SI epithelial layer 30 days following oral administration of OVA. Data summarize two independent experiments. Error bars: means ± s.e.m. Each symbol represents an individual mouse. Statistics were calculated by two-tailed unpaired t-test; *P < 0.05; **P < 0.01; ***P < 0.001.

### pTreg regulate dietary CD8αβ response

Our findings thus far indicate that at steady state, oral antigen-specific T cell responses, in contrast to those triggered by pathogens, do not include an effector component. Inducing effector food-specific CD8αβ T cell responses could potentially have severe consequences, including attack on epithelial cells that present food antigens, as seen in celiac disease^23,24^. Thus we aimed to uncover the mechanisms that restrain effector food-specific CD8αβ responses. Recent studies have demonstrated that Treg cells can guide cDC1 to induce CD8 exhaustion in cancer^25^. Based on this, we hypothesized that under homeostatic conditions cDC1 present dietary antigens and interact with pTreg cells^12^ not to initiate pTreg response, but rather to be educated by food-specific pTreg cells to restrain food-specific CD8αβ responses. To test this, we first analyzed the CD8αβ response to food in MHCII-OFF^ΔRORγt^ mice incapable of generating a pTreg response. Indeed, transferred naive OT-I cells exhibited increased expansion in MHCII-OFF^ΔRORγt^ mice compared to control mice (Fig. 3a,b), suggesting that pTreg cells limit the expansion of food-specific CD8αβ responses.

**Figure 3:**
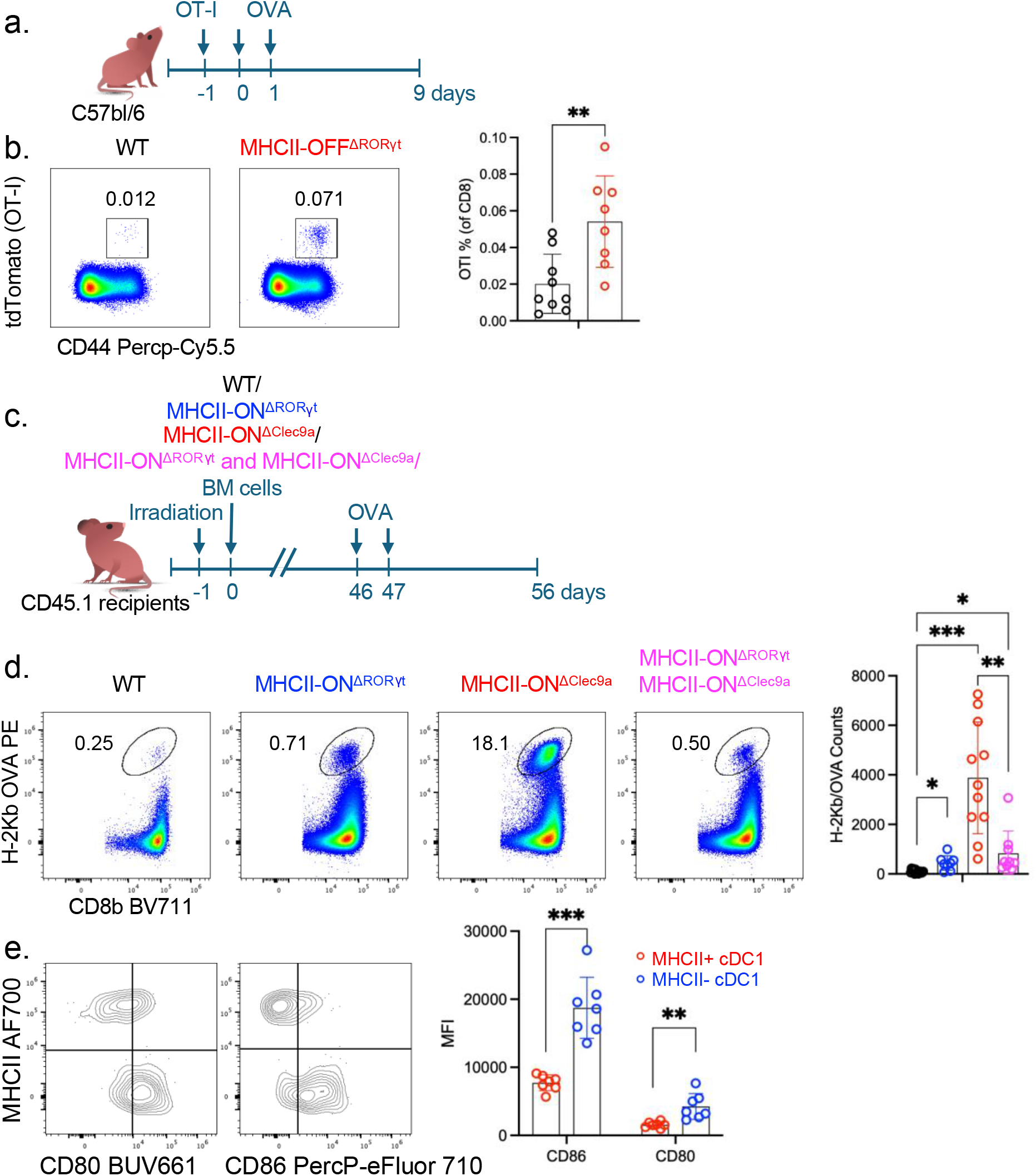
pTreg interact with cDC1 to restrict CD8αβ response. (a) Experimental design of testing food-specific CD8αβ response in MHCII-OFF^ΔRORγt^ mice, lacking food-specific pTreg response. (b) Representative flow cytometry (left) and aggregate data (right) of of OT-I cells in the MLN of MHCII-OFF^ΔRORγt^ (n = 8) and littermate control (MHCII-/fl)(n=9) that were orally administrated with OVA, 10 days after transfer of naive TCR transgenic T cells. Statistics were calculated by two-tailed unpaired t-test; (c) Experimental setup for studying food-specific CD8αβ response in BM chimeric mice, reconstitute with alternative BM cells allowing exclusive antigen presentation by APC subsets, as indicated. (d) Representative flow cytometry profiles (left) and aggregate data (right) of OVA-specific CD8 T cell in the SILP of BM chimeric mice reconstituted with donor bone marrow cells as indicated, that were orally administrated with OVA, 10 days following OVA administration. wild-type (WT) (n = 20), MHCII-ON^ΔRORγt^ (n = 13), MHCII-ON^ΔClec9a^ (n = 18) and MHCII-ON^ΔRORγt^ and MHCII-ON^ΔClec9a^ (n = 15). Data summarize two independent experiments. Error bars: means ± s.e.m. Each symbol represents an individual mouse. (e) representative flow cytometry profiles (left) and aggregate data (right) of CD80 and CD86 expression on MHCII sufficient or deficient cDC1 isolated from MLN of BM chimeric mice reconstituted with a mix of MHCII-ON^ΔRORγt^ and MHCII-ON^ΔClec9a^ donor cells (n=7). cDC1 were gated as dump- (CD19-TCRβ−), CD11c+, PDL2+, CD11b-, CD103+. Error bars: means ± s.e.m. Each symbol represents an individual mouse. Statistics in b and d were calculated by two-tailed unpaired t-test and in e by two-tailed paired t-test ; *P < 0.05; **P < 0.01; ***P < 0.001.

We additionally tested the expansion of food-specific CD8 cells in chimeric mice reconstituted with different combinations of donor cells restricting antigen presentation to specific cell subsets (Fig. 3c). While a minimal CD73+ FR4+ food-specific CD8 response was observed in chimeric mice reconstituted with WT donor cells (Fig. 3d and Extended Data Fig. 3b), mice reconstituted with MHCII-ON^ΔRORγt^ donor cells, which lack MHCII+ cDC1, had an increase in OVA-specific CD8 T cells (Fig. 3d). Moreover, mice reconstituted with MHCII-ON^ΔClec9a^ donor cells, which are deprived of pTreg responses but have OVA-specific Th1 cells, demonstrated an even greater expansion of food-specific CD8 T cells (Fig. 3d), In addition, in this group, food-specific CD8 cells, exhibited reduced expression of FR4 and CD73, along with an elevation of the KLRG1 effector marker (Extended Data Fig. 3b). These findings strongly suggest that pTreg limit the expansion of food-specific CD8 responses, by creating immune synapse interactions with cDC1. They further suggest that effector CD4 T cells, potentially Th1 cells ^16^ (Extended Data Fig. 1c), may further enhance food-specific CD8αβ expansion and instruct a CD8 effector program. Remarkably, chimeric mice reconstituted with a mixture of MHCII-ON^ΔRORγt^ and MHCII-ON^ΔClec9a^ donor cells, thereby generating a pTreg response capable of interacting with cDC1, exhibited a restricted dietary CD8 response (Fig. 3d).

One of the means by which Treg cells suppress the expansion of effector T cells is through the depletion of CD80/CD86 costimulatory molecules from APCs by trogocytosis^26–28^. We therefore analysed cDC1 cells sufficient or deficient for MHCII, isolated from chimeric mice reconstituted with a mix of MHCII-ON ^ΔRORγt^ and MHCII-ON^ΔClec9a^ BM cells and measured CD80 and CD86 expression. MHCII-sufficient cDC1 exhibited decreased levels of both CD80 and CD86 (Fig. 3e), suggesting that pTreg cells may restrict CD8 expansion by depleting CD80/CD86, potentially via trogocytosis. Together, our data suggest that specific antigen-presenting cellular circuits orchestrate the generation of immune responses to food antigens.

### Effector food-specific CD8αβ cells in infections

The mucosal barrier must simultaneously tolerate dietary antigens and guard against opportunistic pathogens. Pathogens mimicking or contaminating food challenge the immune system, which needs to swiftly eliminate them and restore tolerance to dietary antigens to prevent food sensitivities once the threat is resolved ^24^. The mechanisms by which the immune system manages such ‘Trojan horse’ attacks, especially considering immunosuppression by pTregs, and whether that kind of defense results in loss of tolerance to dietary antigens^29^, remain largely unknown.

To address this, we leveraged a well-known intestinal pathogen, *Listeria monocytogenes*, that stably expresses OVA (Lm-OVA) and characterized the immune response in the presence and absence of prior exposure to oral OVA (Fig. 4a). In mice infected with Lm-OVA, OVA-specific CD4 T cells acquired a Th1 program (Fig. 4b,c); by comparison, oral administration of OVA prior to infection resulted in inhibition of the Th1 program (Fig. 4b,c), aligning with the anticipated immunosuppressive function of pTreg cells. Unlike CD4 T-cell responses, prior oral administration of OVA did not prevent endogenous and OT-I CD8αβ cells from acquiring effector markers following infection with Lm-OVA (Fig. 4d-g, Extended Data Fig. 4a,b). In fact, mice that received dietary OVA exhibited enhanced expansion of endogenous OVA-specific CD8 effector T cells upon Lm-OVA infection (Fig. 4d,e), suggesting that in the event of a ‘Trojan horse’ threat, CD8αβ T cells may provide heterologous protection.

**Figure 4:**
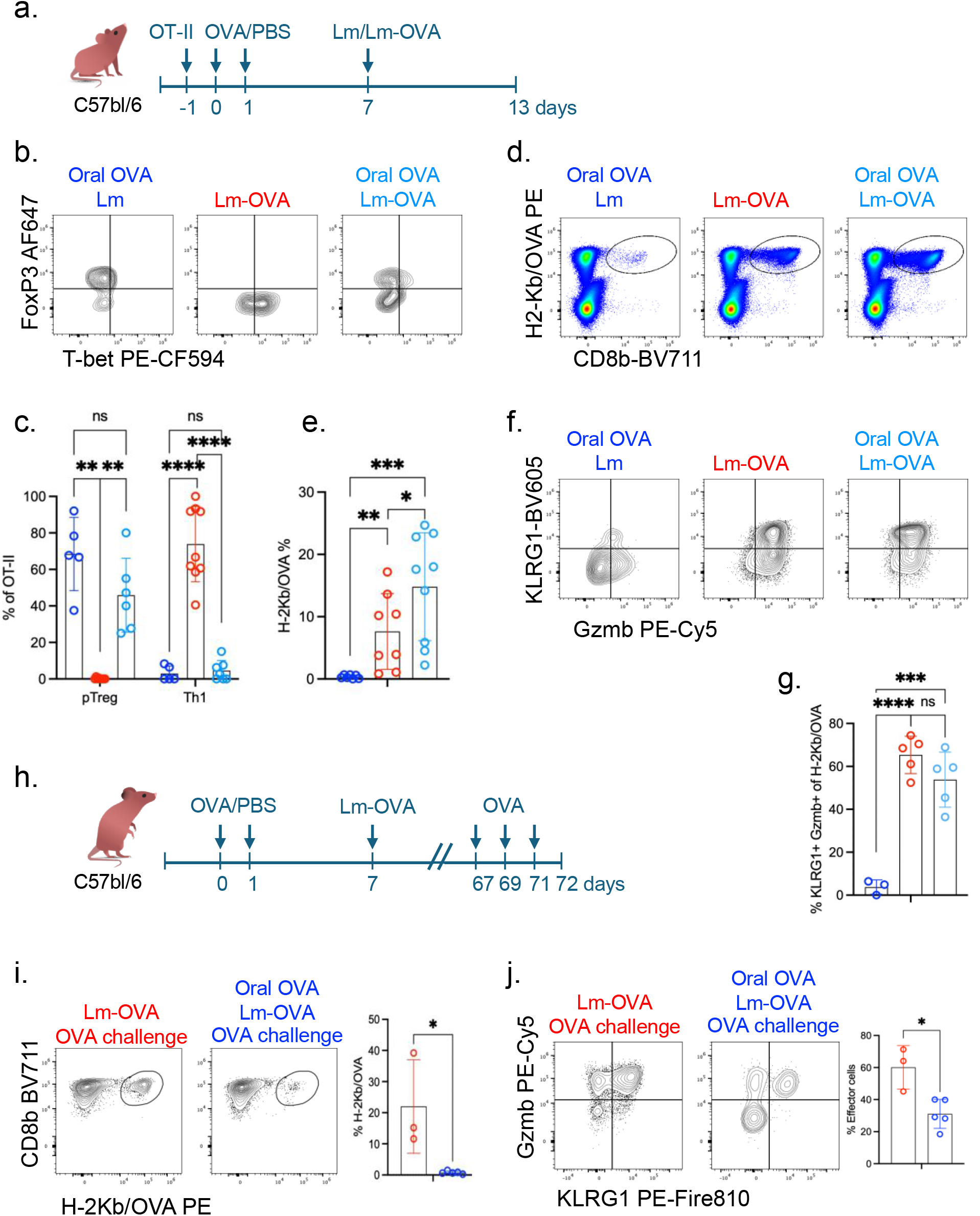
CD8**αβ** response to food antigen in infections. (a) Experimental design to test impact of immune responses to dietary antigens on protective immune responses against pathogens. (b, c) 7 days following naïve OT-II transfer and exposure to OVA or PBS orally, mice were infected with Lm or with Lm-OVA as indicated. 6 days following infection, OT-II CD4 cells (b,c) and endogenous OVA-specific CD8 T cells (d-g) from the SILP were analyzed. Representative flow cytometry profiles (b) and aggregate data (c) of OT-II differentiation. Oral OVA and Lm (n = 5), Lm-OVA (n = 9), Oral OVA and Lm-OVA (n = 6). Representative flow cytometry profiles (d,f) and aggregate data (e,g) of OVA-specific CD8 effector expansion and effector program. Oral OVA and Lm (n = 3), Lm-OVA (n = 5), Oral OVA and Lm-OVA (n = 5). Data in b-g summarize two or three independent experiments. Error bars: means ± s.e.m. Each symbol represents an individual mouse. Statistics were calculated by two-tailed unpaired t-test; (h) Experimental setup to study the long-lasting implcations of immune responses to dietary antigens. 7 days following naïve OT-II transfer and exposure to OVA or PBS orally, mice were infected with Lm-OVA as indicated. 45 days following infection, mice were re-challenged with OVA orally. (i,j) Representative flow cytometry profiles (left) and aggregate data (right) demonstrating the expansion of endogenous OVA-specific CD8αβ in the SILP (i) and their effector program (j). Statistics was calculated by a non-parametic Mann-Whitney U test. Error bars: means ± s.e.m. Each symbol represents an individual mouse. *P < 0.05; **P < 0.01; ***P < 0.001.

To investigate the prolonged impact of an infection by a pathogen that mimics food, and the host’s ability to regain tolerance, we re-challenged mice with dietary OVA following the resolution of Lm-OVA infection. (Fig. 4h). In mice that had not been exposed to dietary OVA before Lm-OVA infection, an oral re-challenge with OVA led to a marked increase in OVA-specific effector CD8αβ cells (Fig. 4i, j), indicative of developing sensitivity to OVA as food. Conversely, mice that were exposed to dietary OVA before Lm-OVA infection did not show any expansion of cells following an OVA rechallenge (Fig. 4i,j). Thus, food-specific CD8αβ cells retained responsiveness to tolerogenic signals established by exposure to dietary antigens, even following a protective inflammatory response to a pathogen sharing antigenicity with the tolerizing agent. This property of the CD8 T cells sets them apart from the conventional pathogen-induced CD8αβ Trm response^30^. These findings demonstrate that in response to dietary antigens, the immune system activates a ‘Trm-like’ CD8αβ response, which is capable of adopting effector functions that can potentially protect the host from pathogens, while also maintaining tolerance to dietary antigens once the threat has been neutralized. The mechanism by which food-specific CD8 T cells are unleashed from pTreg regulation when there is a threat will require further investigation.

Collectively, these findings illuminate the distinct roles of RORγt-lineage APCs and cDC1s in orchestrating dynamic, balanced immune responses to dietary antigens. They also unveil a complex multicellular network that works in concert to foster an immune response that offers protection within the general strategy of tolerance (Extended Data Fig. 5).

## Discussion

All living organisms must balance the need to absorb nutrients from their environment and allow mutualistic relations with gut microbes while protecting themselves from pathogens. To prevent chronic inflammation induced by dietary antigens and gut commensals, our immune system can generate an anti-inflammatory pTreg response that effectively suppresses CD4+ T cell effector programs. However, pTreg responses to food antigens or to gut microbes may leave the host vulnerable to attacks by pathogens that share these antigens. Alternatively, activating a protective immune response against dietary antigens presented by a pathogen might reduce tolerance toward these food antigens or commensals, potentially leading to adverse reactions during subsequent food consumption and to chronic intestinal inflammation. Here we found that the immune response to food enables acute effector responses within the broader strategy of tolerance. Our data points towards a mechanism that counters pTreg-induced immunosuppression by initiating a protective CD8αβ response. This response can potentially shield the host from ‘Trojan horse’ attacks and, at the same time, unlike traditional immune responses to pathogens in which CD8 Trm cells are rapidly activated upon antigen exposure, it could reinstate tolerance to food antigen, remaining inactive during subsequent encounters with them.

Past work suggesting that cDC1s present dietary antigens and interact with food-specific CD4 T cells, together with experiments failing to show loss of food-specific pTreg cells or loss of oral tolerance in mice lacking the cDC1 subset, has contributed to a widely held view that DC subsets are redundant in their ability to induce pTreg responses. Here, in contrast, we show that RORγt+ APCs exclusively initiate the pTreg response to dietary antigens in non-redundant fashion, and that following differentiation these pTreg cells interact with cDC1s to shape ‘Trm-like’ food-specific CD8αβ responses. These findings reveal that immune responses to food involve a dynamic antigen-specific cellular network comprised of counterbalancing T cell subsets. Understanding the contributions, composition, regulation, and mechanism of this cellular network may shed light on inflammatory conditions including allergy and celiac disease and suggest novel therapeutic targets.

## Methods

### Mice

Mice were bred and maintained in the animal facility of the Weizmann Institute, in specific pathogen-free conditions. C57BL/6 mice (Jax 000664), Itgav f/f (B6.129P2(Cg)-Itgav tm2Hyn/J Jax 032297, CD45.1 mice (B6.SJL-Ptprca Pepcb/BoyJ, Jax 002014), I-AB f/f (B6.129X1-H2-Ab1tm1Koni/J Jax 013181), OT-II (B6.Cg-Tg(TcraTcrb)425Cbn/J, Jax 004194), OT-I (C57BL/6-Tg(TcraTcrb)1100Mjb/J, Jax 003831) and CD90.1 (B6.PL-Thy1a/CyJ Jax 000406) mice were purchased from Jackson Laboratories. RORγt-cre mice^31^ were generated by Dan R. Littman’s lab, I-AB lox-STOP-lox were generated by Terri M. Laufer’s lab^32^, and Clec9a-cre^33^ mice were generated by Caetano Reis e Sousa’s lab. Littermates with matched sex (both males and females) were used. Mice in all the experiments were 6–12 weeks old at the starting point of treatment. Animal sample size estimates were determined using power analysis (power = 90% and α = 0.05) based on the mean and s.d. from our previous studies and/or pilot studies using 4–5 animals per group. All animal procedures were performed in accordance with protocols approved by the Institutional Animal Care and Usage Committee of the Weizmann Institute.

### Antibodies, intracellular staining and flow cytometry

The following monoclonal antibodies were purchased from eBiosciences, BD Pharmingen or BioLegend: PE/Cyanine5 Granzyme B (clone QA16A02), Alexa Fluor® 700 I-A/I-E (clone M5/114.15.2), FITC TCR Vβ5.1, 5.2 (clone MR9-4), Alexa Fluor® 594 TCR β chain (clone H57-597), Brilliant Violet 711™ CD8b (Ly-3) (clone YTS156.7.7), PE/Cyanine7 FR4 (Folate Receptor 4) (clone 12A5), CD25 Spark-NIR 685 (clone PC61), CD11c PE-Cy7 (clone N418), CD73 Pacific Blue (clone TY/11.8), CD69 FITC (clone H1.2F3), KLRG1 PE-Fire810 (clone 2F1/KLRG1), CD107a (LAMP-1) APC-Cy7 (clone 1D4B), CD51 PE (clone RMV-7), PD-L2 (CD273, B7-DC) APC (clone TY25), XCR1 APC-Cy7 (clone ZET).

BD Bioscience: PE-CF594 T-BET (clone O4–46), RB705 CD80 (clone 16-10A1), CD45.1 BV650 (clone A20), CD11c BUV496 (clone HL3), CD103 BV510 (clone M290), CD25 PE (clone PC61), GATA3 BUV395 (clone L50-823), CD86 BUV661 (clone PO3), Bcl6 BUV805 (clone K112-91), CD4 APC-Cy7 (clone GK1.5), CD45.2 BUV737 (clone 104), CD44 Percp-Cy5.5 (clone IM7), RORgt BV421 (clone Q31-378), FoxP3 AF647 (clone R16-715), CD19 RB545 (clone 1D3), CD90.1 BV786 (clone OX-7), SIRPa (CD172a) BUV615 (clone P84), CCR9 (CD199) BV421 (clone CW-1.2), CD11b BUV805 (clone M1/70), CXCR6 (clone CD186) APC (clone SA051D1), CD62L BV421 (clone MEL-14), Clec12A (CD371) PE (clone 5D3),

T-Select H-2Kb OVA Tetramer-SIINFEKL-PE (MBL), H-2K(b) chicken ova 257-264 SIINFEKL (NIH Tetramer Core Facility). Live/dead Fixable Viability Stain 575V (BD Horizon) was used to exclude dead cells.

For transcription factor staining, cells were stained for surface markers, followed by fixation and permeabilization before nuclear factor staining according to the manufacturer’s protocol (FOXP3 staining buffer set from eBioscience). Flow cytometric analysis was performed on an Cytek Aurora (Cytek) and analysed using FlowJo 10.8.1 software (Tree Star).

### Flow cytometry gating strategy

OT-II gating: FSC, SSC; Live Dead−, singlets, Dump− (B220, MHCII), TCRβ+, CD8−, CD4+, VB5+, CD45.1+ or CD90.1+; OT-I gating: FSC, SSC; Live Dead−, singlets, Dump− (B220, MHCII), TCRβ+, CD4−, CD8+, VB5+, CD45.1+ or CD90.1+ or tdTomato+; H-2Kb/OVA gating: FSC, SSC; Live Dead−, singlets, Dump− (B220, MHCII), TCRβ+, CD4−, CD8+, H-2Kb/OVA+.

### Isolation of lymphocytes and APCs

After removal of Peyer’s patches, small intestine tissues were treated sequentially with PBS containing 1 mM DTT at room temperature for 10 min, twice with 5 mM EDTA at 37 °C for 10 min to remove epithelial cells, and then minced and dissociated in digestion buffer (RPMI containing collagenase (1 mg ml−1 collagenase D; Roche), DNase I (100 μg ml−1; Sigma), dispase (0.1 U ml−1; Worthington) and 10% FBS) with constant stirring at 37 °C 55 min. Leukocytes were collected at the interface of a 40%–80% Percoll gradient (GE Healthcare). Lymph nodes were mechanically disrupted for lymphocyte isolation. For isolation of myeloid cells and ILC, lymph nodes were mechanically disrupted with digestion buffer with constant stirring at 37 °C 30 min.

### Oral administration of OVA

OVA (grade III, Sigma, A5378) was administered intragastrically using plastic gavage needles. Mice were gavaged with 50mg in 200µl of PBS, 1 and 2 days after adoptive transfer of OT-II or OT-I T cells. In some experiments, as indicated, mice were simultaneously fed ad libitum with OVA dissolved in drinking water (1.5% w/v).

### Infection with Listeria and Lm-OVA

For infection experiments, ovalbumin-expressing Listeria (Lm-OVA). (PMID 11207297) or the parental wild-type stain 10403S (Lm) were used. Mice were infected orally with 10^9 CFU Lm-OVA or Listeria, 7 days following oral administration of OVA.

### Adoptive transfer of OT-II and OT-I TCR transgenic cells

Adoptive transfer of OT-II and OT-I was done as was previously described, with minor modifications. Spleens and lymph nodes from donor OT-II and OT-I TCR transgenic mice were collected and mechanically disassociated. Red blood cells were lysed using ACK lysis buffer (Lonza). Naive OT-II T cells were sorted as CD4+TCRβ+CD44loCD62LhiCD25−Vβ5+CD45.1+, or as CD4+TCRβ+CD44loCD62LhiCD25−Vβ5+CD90.1+, Naive OT-I T cells were sorted as CD8+TCRβ+CD44loCD62LhiCD25−Vβ5+tdTomato+, or as CD8+TCRβ+CD44loCD62LhiCD25−Vβ5+CD45.1+ on the MA900 Multi-Application Cell Sorter (SONY). For analysis of early differentiation, cells were additionally labelled with CFSE (Thermo Fisher) or with cell proliferation V450 (BD Bioscience). Cells were resuspended in PBS on ice and 50,000-100,000 cells were then transferred into congenic isotype-labelled recipient mice by retro-orbital injection. Cells from MLN were analysed 3 days after transfer and cells from small intestine LP were analyzed 9–14 days after transfer.

### Generation of bone marrow chimeric reconstituted mice

To generate chimeric mice, 4-to 5-week-old CD45.1 mice were irradiated twice with 500 rads per mouse at an interval of 2–5 h (X-RAD 320 X-Ray Irradiator). One day later, bone marrow mononuclear cells were isolated from donor mice, as indicated in each experiment, by flushing the femur bones. Red blood cells were lysed with ACK lysing buffer, and lymphocytes were depleted for Thy1.2 using magnetic microbeads (Miltenyi). Bone marrow cells were resuspended in PBS and a total 2 to 5 × 10^6^ BM cells were injected intravenously into the irradiated mice. In case of mixed bone marrow cell reconstitution, a ratio of 1:1 was used. Mice were kept for a week on broad spectrum antibiotics (1 mg ml−1 sulfamethoxazole and 0.2 mg/ml−1 trimethoprim), followed by microbiome reconstitution by fecal gavage. Mice were reconstituted for 1–2 months before OVA administration. After 7 weeks, peripheral blood samples were collected and analysed by FACS to check for reconstitution.

### Statistical analysis

For animal studies, mutant and control groups did not always have similar s.d. values and therefore an unpaired two-sided Welch’s t-test was used. Error bars represent ± s.d. Mouse sample size estimates were determined based on our previous studies and/or pilot studies using 4–5 mice. No samples were excluded from analysis. P < 0.05 was considered significant: * p < 0.05; ** p < 0.01; ***p < 0.001; ****p < 0.0001. Details as to number of replicates, sample size, significance tests, and value and meaning of n for each experiment are included in the Methods or Figure legends.

## Acknowledgements

We thank members of the Kedmi laboratory, D. R. Littman, J. M. Gardner, J. Talbot K. R. Mesa O. J. Harrison and S. Naik, for valuable discussion and critical reading of the manuscript; R. Dahan for sharing mice strains; B. Sheridan for advice and providing Lm-OVA; Weizmann transgenic core unit for rederivation of mutant mice. This work was supported by the Abisch-Frenkel RNA Therapeutics Center and by a research grant from the Weizmann SABRA - Yeda-Sela - WRC Program, the Estate of Emile Mimran, and The Maurice and Vivienne Wohl Biology Endowment.

## Contributions

A.R. and R.K. designed the study and analysed the data. A.R., H.O and J.T. performed mouse genetic experiments. R.K. wrote the manuscript, with input from the other authors and supervised the research.

