## Supplementary Data for "Coordinated network of T cells and antigen presenting cells regulate tolerance to food"

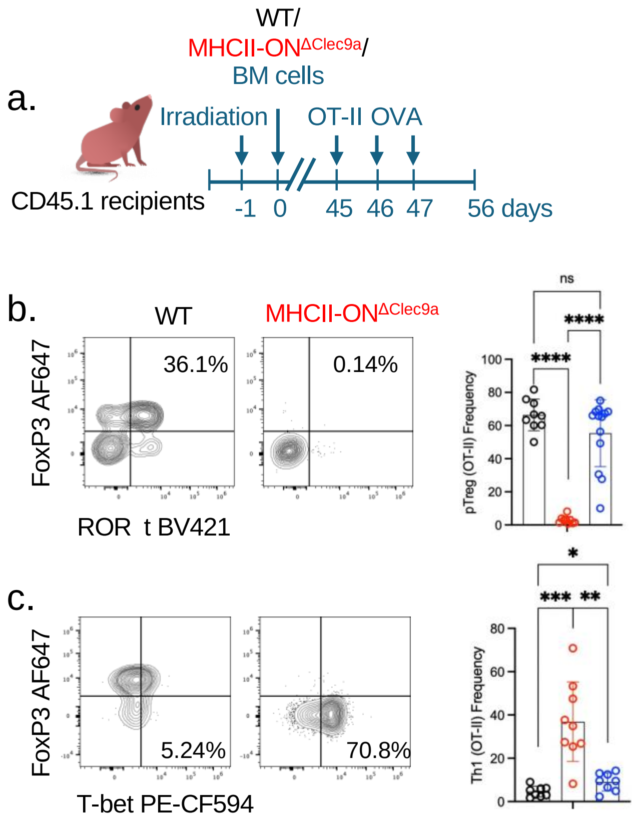


Extended Data Figure 1: **RORγt APCs and not cDC1 induces pTreg programs in response to dietary antigens**. (a) Experimental design of studying food-specific pTreg induction in BM chimeric mice, reconstitute with alternative BM cells allowing exclusive antigen presentation by APC subsets, as indicated. (b) Representative flow cytometry profiles (left) and aggregate data (right) of OT-II T cell differentiation in SILP of bone marrow chimeric mice reconstituted with donor bone marrow cells as indicated, that were orally administrated with OVA, 10 days after transfer of naive TCR transgenic T cells. wild-type (WT) (n = 9), MHCII-ON$^{\Delta Clec9a}$ (n = 9), MHCII-ON$^{\Delta ROR\gamma t}$ and MHCII-ON$^{\Delta Clec9a}$ (n = 10) (c) Data summarize three independent experiments. Error bars: means ± s.e.m. Each symbol represents an individual mouse. Statistics were calculated by two-tailed unpaired t-test; *$P < 0.05$; **$P < 0.01$; ***$P < 0.001$.

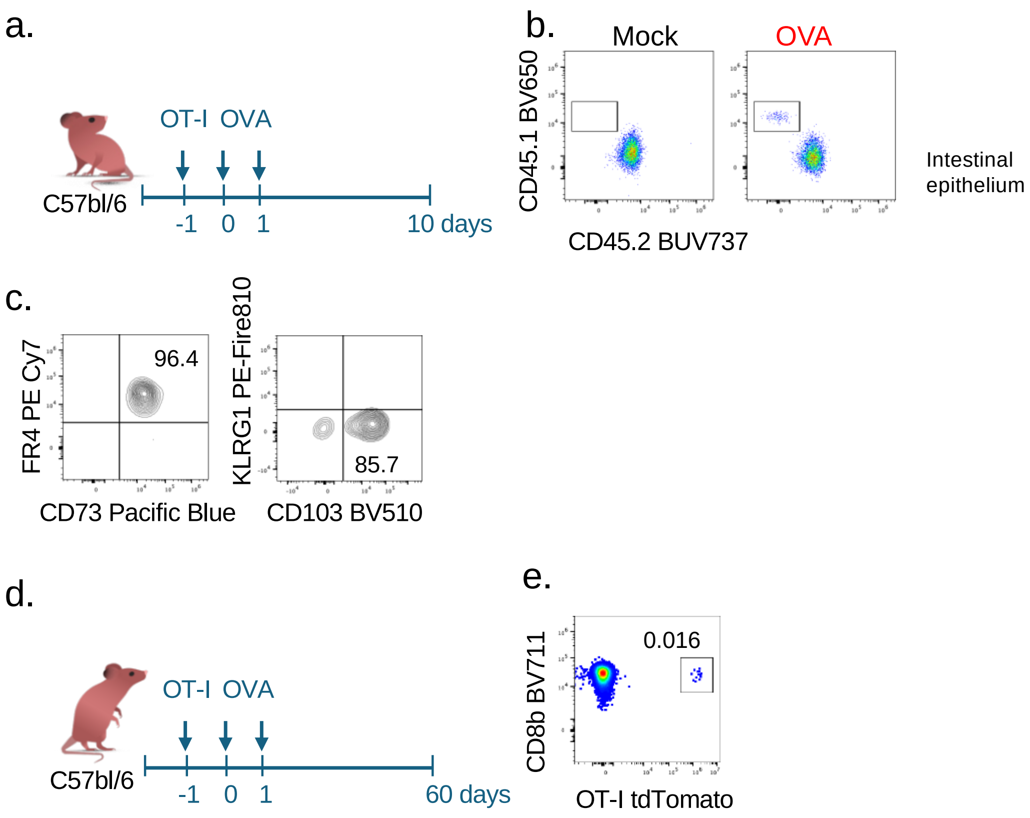


Extended Data Figure 2: **cDC1 induces a food-specific intra-epithelial lymphocytes 'Trm-like' CD8αβ response** (a,d) Experimental design of testing food-specific CD8 response in the SI epithelial layer (b,c) Representative flow cytometry profiles of OT-I expansion in the SI epithelial layer in mice orally administrated with OVA or PBS, 10 days following naïve OT-I transfer. Data summarize two independent experiments. (e) Representative flow cytometry profiles of OT-I expansion in the SILP in mice orally administrated with OVA, 60 days following naïve OT-I transfer.

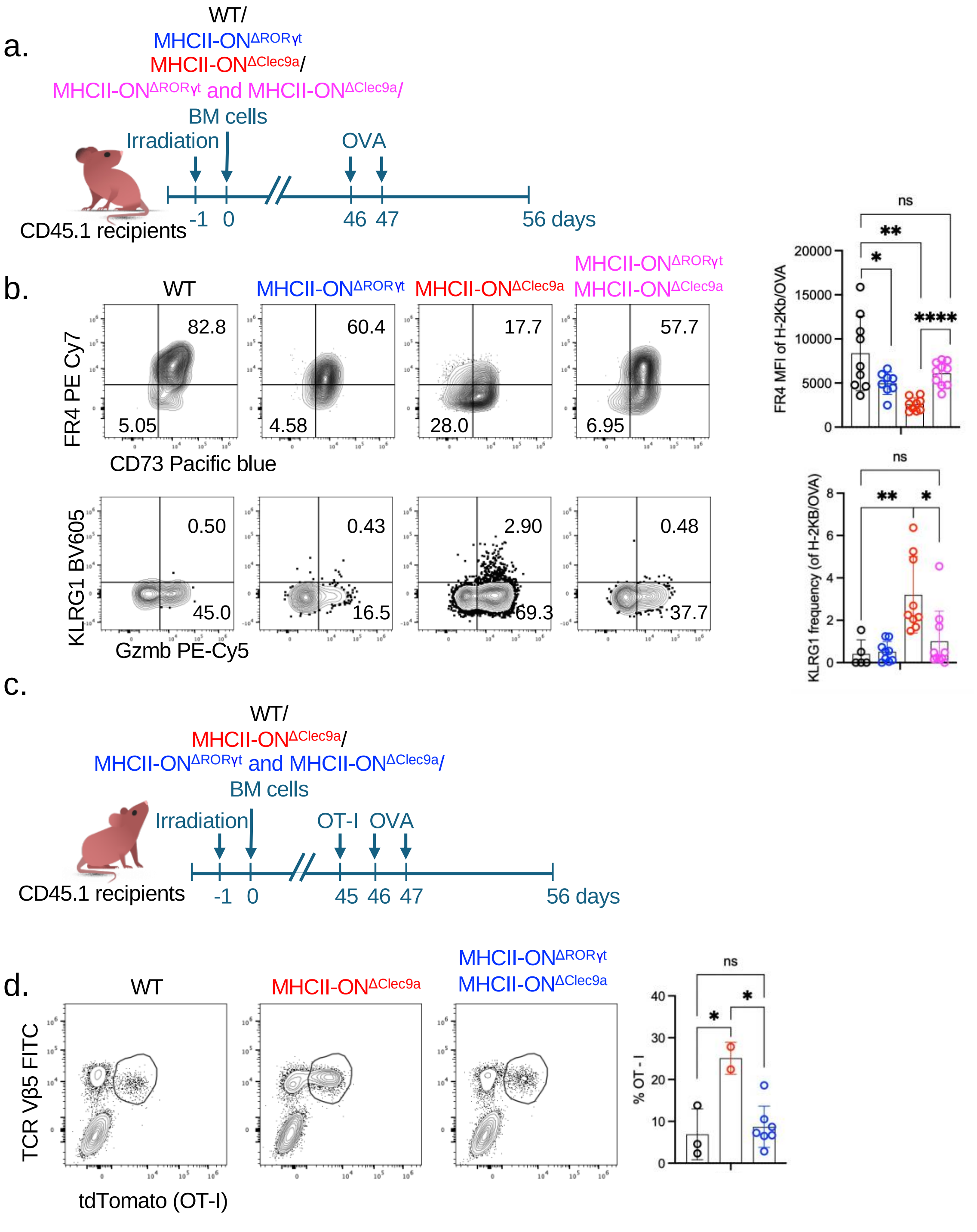


Extended Data Figure 3: **pTreg instruct cDC1 and shape CD8αβ response**. (a) Experimental setup for studying food-specific CD8αβ response in BM chimeric mice, reconstitute with alternative BM cells allowing exclusive antigen presentation by APC subsets, as indicated. (b) Representative flow cytometry profiles (left) and aggregate data (right) presenting FR4 and CD73 markers (upper panel) or Gzmb and KLRG1 (lower pannel) in OVA-specific CD8 T cell in the SILP of BM chimeric mice reconstituted with donor bone marrow cells as indicated, that were orally administrated with OVA, 10 days following OVA administration. wild-type (WT) (n = 9), MHCII-ON$^{\Delta ROR\gamma t}$ (n = 8), MHCII-ON$^{\Delta Clec9a}$ (n = 9) and MHCII-

ON$^{\Delta ROR\gamma t}$ and MHCII-ON$^{\Delta Clec9a}$ (n = 10). Data summarize three independent experiments. Error bars: means ± s.e.m. Each symbol represents an individual mouse. (c) Experimental setup for studying food-specific CD8αβ response in BM chimeric mice, reconstitute with alternative BM cells allowing exclusive antigen presentation by APC subsets, as indicated. (d) Representative flow cytometry profiles (left) and aggregate data (right) of OT-I cells in the SILP of BM chimeric mice reconstituted with donor bone marrow cells as indicated, that were orally administrated with OVA, 12 days following OVA administration. wild-type (WT) (n = 3), MHCII-ON$^{\Delta Clec9a}$ (n = 2) and MHCII-ON$^{\Delta ROR\gamma t}$ and MHCII-ON$^{\Delta Clec9a}$ (n = 7). Data summarize two independent experiments. Error bars: means ± s.e.m. Each symbol represents an individual mouse. Statistics were calculated by two-tailed unpaired t-test; *$P < 0.05$; **$P < 0.01$; ***$P < 0.001$.

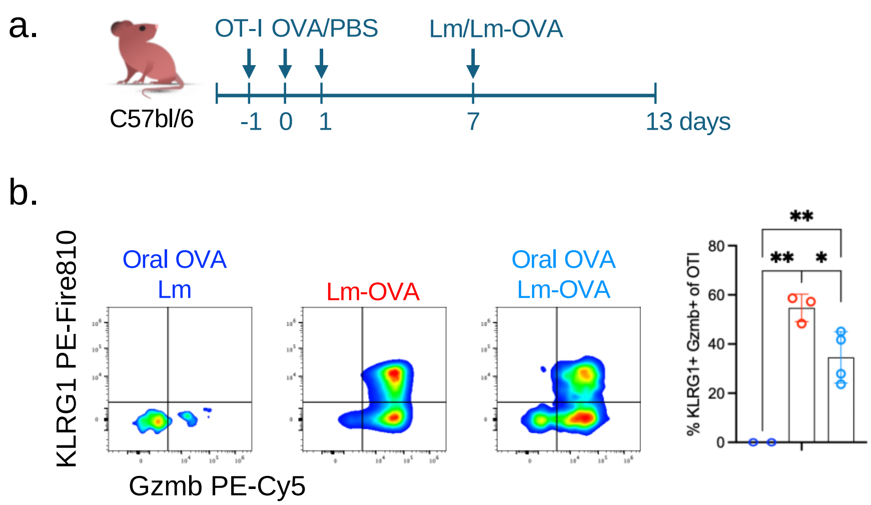


Extended Data Figure 4: CD8**αβ** response to food antigen in infections.
(a) 7 days following naïve OT-I transfer and exposure to OVA or PBS orally, mice were infected with Lm or with Lm-OVA as indicated. 6 days following infection, OT-I cells from the SILP were analyzed. Representative flow cytometry profiles (left) and aggregate data (right) of OT-I effector program. Error bars: means ± s.e.m. Each symbol represents an individual mouse. Statistics were calculated by two-tailed unpaired t-test; *P < 0.05; **P < 0.01; ***P < 0.001.

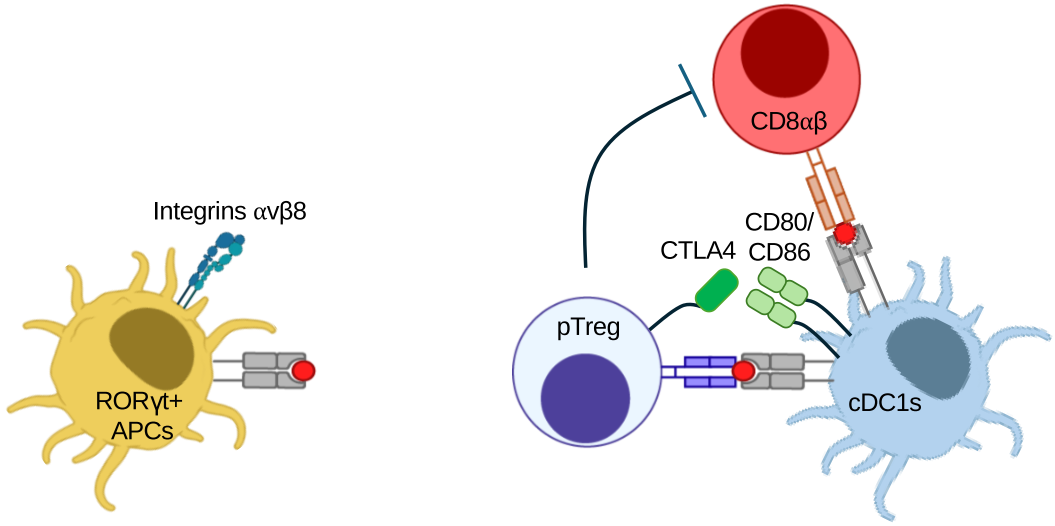


Extended Data Figure 5: **Schematics of the immune response to dietary antigens, which is generated by a dynamic, antigen-specific cellular network**. Antigen presentation and integrin $\alpha_v\beta_8$ are required in RORγt$^+$ APCs for pTreg cell differentiation in response to food antigens. Following their differentiation pTregs interact with cDC1and reduce the expression of CD80/CD86 by trogocytosis to restrict CD8αβ response to food.
